# Reconstructing Noisy Gene Regulation Dynamics Using Extrinsic-Noise-Driven Neural Stochastic Differential Equations

**DOI:** 10.1101/2025.03.11.642678

**Authors:** Jiancheng Zhang, Xiangting Li, Xiaolu Guo, Zhaoyi You, Lucas Böttcher, Alex Mogilner, Alexander Hoffmann, Tom Chou, Mingtao Xia

**Affiliations:** Department of Electrical and Computer Engineering, University of California, Riverside, CA, USA; Department of Computational Medicine, University of California, Los Angeles, CA, USA; Department of Microbiology, Immunology, and Molecular Genetics (MIMG), & Institute for Quantitative and Computational Biosciences, University of California Los Angeles, Los Angeles, CA, USA; Ray and Stephanie Lane Computational Biology Department, School of Computer Science, Carnegie Mellon University, Pittsburgh, PA, USA; Department of Computational Science and Philosophy, Frankfurt School of Finance and Management, Frankfurt am Main, Germany; Laboratory for Systems Medicine, Department of Medicine, University of Florida, FL 32610, USA; Courant Institute of Mathematical Sciences, New York University, NY 10012, USA; Department of Mathematics, University of California, Los Angeles, CA 90095, USA; Department of Mathematics, University of Houston, Houston, TX 77004, USA

**Author notes:** (indicates equal contribution) (Jiancheng Zhang, Xiangting Li, Xiaolu Guo).

## Abstract

Proper regulation of cell signaling and gene expression is crucial for maintaining cellular function, development, and adaptation to environmental changes. Reaction dynamics in cell populations is often noisy because of (i) inherent stochasticity of intracellular biochemical reactions (“intrinsic noise”) and (ii) heterogeneity of cellular states across different cells that are influenced by external factors (“extrinsic noise”). In this work, we introduce an extrinsic-noise-driven neural stochastic differential equation (END-nSDE) framework that utilizes the Wasserstein distance to accurately reconstruct SDEs from trajectory data from a heterogeneous population of cells (extrinsic noise). We demonstrate the effectiveness of our approach using both simulated and experimental data from three different systems in cell biology: (i) circadian rhythms, (ii) RPA-DNA binding dynamics, and (iii) NF*κ*B signaling process. Our END-nSDE reconstruction method can model how cellular heterogeneity (extrinsic noise) modulates reaction dynamics in the presence of intrinsic noise. It also outperforms existing time-series analysis methods such as recurrent neural networks (RNNs) and long short-term memory networks (LSTMs). By inferring cellular heterogeneities from data, our END-nSDE reconstruction method can reproduce noisy dynamics observed in experiments. In summary, the reconstruction method we propose offers a useful surrogate modeling approach for complex biophysical processes, where high-fidelity mechanistic models may be impractical.

## I. INTRODUCTION

Reactions that control signaling and gene regulation are important for maintaining cellular function, development, and adaptation to environmental changes, which impact all aspects of biological systems, from embryonic development to an organism’s ability to sense and respond to environmental signals. Variations in gene regulation, arising from noisy biochemical processes [1, 2], can result in phenotypic heterogeneity even in a population of genetically identical cells [3].

Noise within cell populations can be categorized as (i) “intrinsic noise,” which arises from the inherent stochasticity of biochemical reactions and quantifies, *e*.*g*., biological variability across cells in the same state [2, 4, 5], and (ii) “extrinsic noise,” which encompasses heterogeneities in environmental factors or differences in cell state across a population. A substantial body of literature has focused on quantifying intrinsic and extrinsic noise from experimental and statistical perspectives [1, 2, 6–13]. Experimental studies have specifically identified relevant sources of noise in various organisms, including *E. coli* (Escherichia coli), yeast, and mammalian systems [2, 14–17].

Extrinsic noise is associated with uncertainties in biological parameters that vary across different cells. The distribution over physical and chemical parameters determine the observed variations in cell states, concentrations, locations of regulatory proteins and polymerases [1, 2, 18], and transcription and translation rates [19]. For example, extrinsic noise is the main contributor to the variability of concentrations of oscillating p53 protein levels across cell populations [20]. On the other hand, intrinsic noise, *i*.*e*., inherent stochasticity of cells in the same state, can limit the accuracy of expression and signal transmission [2, 5]. Based on the law of mass action [21, 22], ordinary differential equations (ODEs) apply only in some deterministic or averaged limit and do not take into account intrinsic noise. Therefore, stochastic models are necessary to accurately represent biological processes, such as thermodynamic fluctuations inherent to molecular interactions within regulatory networks [1, 5, 18] or random event times in birth-death processes.

Existing stochastic modeling methods that account for intrinsic noise include Markov jump processes [23, 24] and SDEs [25–27]. Additionally, a hierarchical Markov model was designed in [28] for parameter inference in dual-reporter experiments to separate the contributions of extrinsic noise, intrinsic noise, and measurement error when both extrinsic and intrinsic noise are present. The described methods have been effective in the reconstruction of low-dimensional noisy biological systems. However, these methods usually require specific forms of a stochastic model with unknown parameters to be inferred. It is unclear whether these methods and their generalizations can be applied to more complex (*e*.*g*., higher-dimensional) systems for which a mechanistic description of the underlying biophysical dynamics is not available or impractical.

In this work, we introduce an extrinsic-noise-driven neural stochastic differential equation (END-nSDE) reconstruction method. Our method builds upon a recently developed Wasserstein distance (*W*_2_ distance) nSDE reconstruction method [29] to identify macromolecular reaction kinetics and cell signaling dynamics from noisy observational data under the presence of both extrinsic and intrinsic noise. A key question we address in this paper is how extrinsic noise that characterizes cellular heterogeneity influences the overall stochastic dynamics of the population. Our approach differs from the one proposed in Ref. [29] in that our method takes into account both cell heterogeneity and intrinsic fluctuations. In Fig. 1, we provide an overview of the systems that we study in this work.

**Fig 1.**
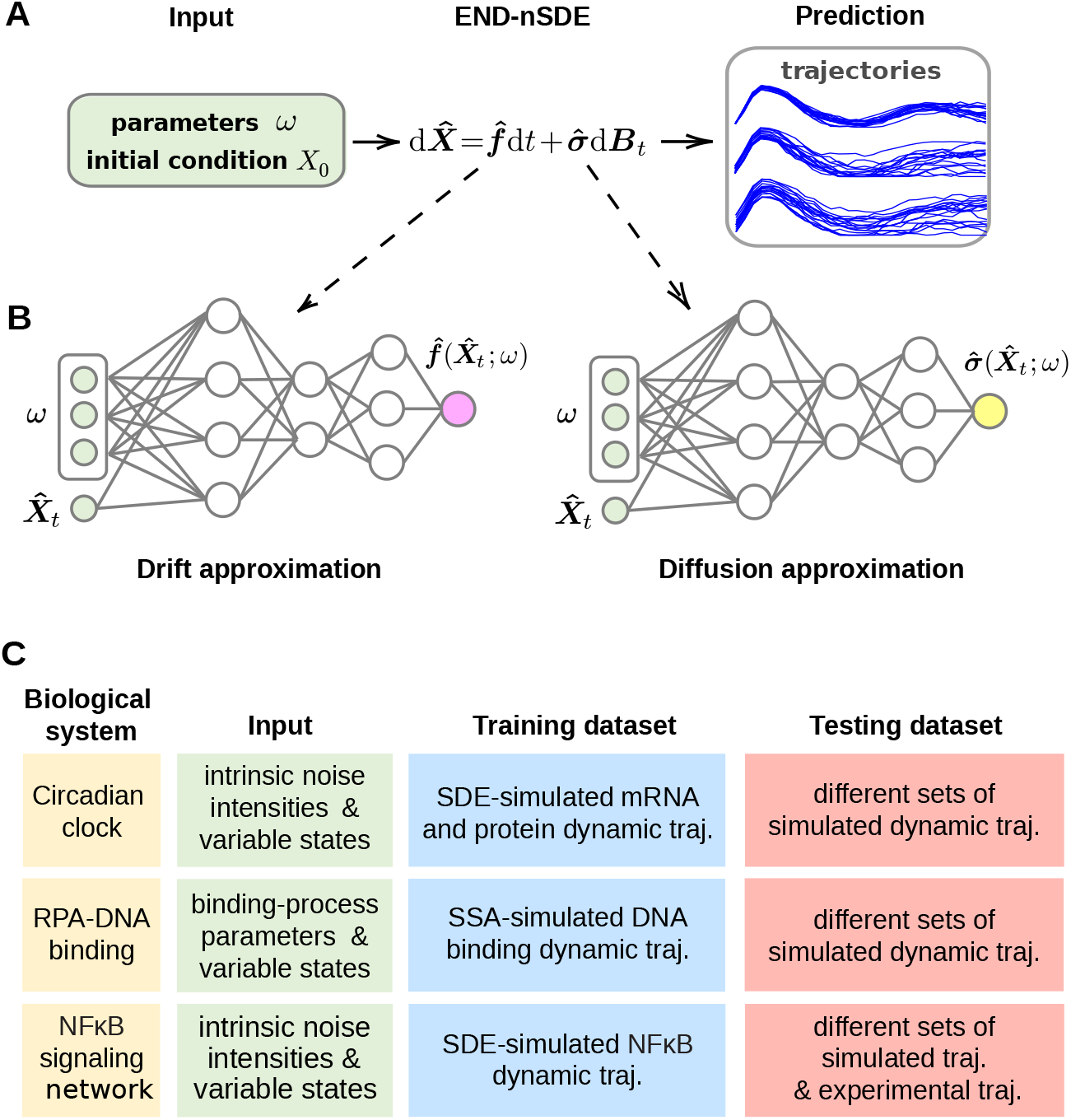
Workflow of our proposed END-nSDE prediction on parameters altering stochastic dynamics. A. The predicted trajectories are generated through the reconstructed SDE 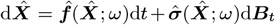.B. The drift and diffusion functions, 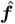 and 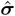,are approximated using parameterized neural networks. The parameterized neural-network-based drift function 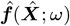 and diffusion function 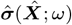 take the system state 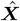 and biological parameters *ω* as inputs. C. Diagram of three examples illustrating the nSDE input, along with training and testing datasets.

Our approach employs neural networks as SDE approximators in conjunction with the `torchsde` package [30, 31] for reconstructing noisy dynamics from data. Previous work showed that for SDE reconstruction tasks, the *W*_2_ distance nSDE reconstruction method outperforms other benchmark methods such as generative adversarial networks [29, 32]. Additionally, the *W*_2_ distance nSDE reconstruction method can directly extract the underlying SDE from temporal trajectories without requiring specific mathematical forms of the terms in the underlying SDE model. We apply our END-nSDE methodology to three biological processes that illustrate (i) circadian clock rhythm, (ii) RPA-DNA binding dynamics, and (iii) NF*κ*B signaling to show that END-nSDE can predict how extrinsic noise modulates stochastic dynamics with intrinsic noise. Additionally, our method demonstrates superior performance compared to several time-series modeling methods including recurrent neural networks (RNNs), long short-term memory networks (LSTMs), and Gaussian processes. In summary, the reconstruction method we propose provides a useful surrogate modeling approach for complex biomedical processes, especially in scenarios where high-fidelity mechanistic models are impractical.

## II. METHODS AND MODELS

In this work, we extend the temporally decoupled squared *W*_2_-distance SDE reconstruction method proposed in Refs. [29, 33] to reconstruct noisy dynamics across a heterogeneous cell population (“extrinsic noise”). Our goal is to not only reconstruct SDEs for approximating noisy cellular signaling dynamics from time-series experimental data, but to also quantify how heterogeneous biological parameters, such as enzyme- or kinase-mediated biochemical reaction rates, affect such noisy cellular signaling dynamics.

### A. SDE reconstruction with heterogeneities in biological parameters

The *W*_2_-distance-based neural SDE reconstruction method proposed in Ref. [29] aims to approximate the SDE

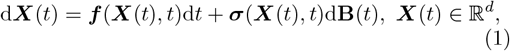

using an approximated SDE

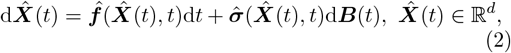

where 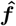 and 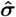 are two parameterized neural networks that approximate the drift and diffusion functions ***f*** and ***σ*** in Eq. (1), respectively. These two neural networks are trained by minimizing a temporally decoupled squared *W*_2_-distance loss function

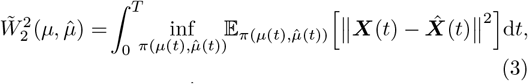

where ***X***(*t*) and 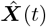 are the observed trajectories at time *t* and trajectories generated by the approximate SDE model Eq. (2) at time *t*, respectively. 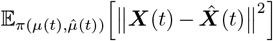 represents the expectation when 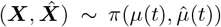.Our *W*_2_-distance-based SDE reconstruction method can result in very small errors 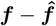 and 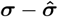 in the reconstructed diffusion and jump functions. The 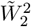 term in Eq. (3) is denoted as the squared temporally decoupled squared *W*_2_ distance loss function. For simplicity, in this paper, we shall also denote the Eq. (3) as the squared *W*_2_ loss. *µ* and 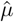 are the probability distributions associated with the stochastic processes {***X***(*t*)}, 0≤ *t* ≤*T* and 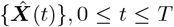, respectively, while *µ*(*t*) and 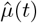 are the probability distributions of ***X***(*t*) and 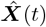 at a specific time *t*. The coupling distributions *π* of two distributions 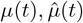 on the probability space ℝ ^*d*^ are defined by

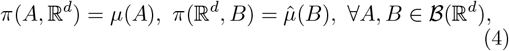

where ℬ(ℝ^*d*^) is the Borel *σ*-algebra on ℝ^*d*^. The infimum in Eq. (3) is taken over all possible coupling distributions 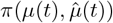 and ∥ *·* ∥ denotes the ℓ^2^ norm of a vector. That is,

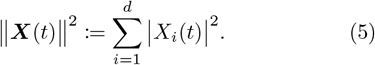

Across different cells, extrinsic noise or cellular heterogeneities such as differences in kinase or enzyme abundances resulting from cellular variabilities, can lead to variable, cell-specific, gene regulatory dynamics. Such heterogeneous and stochastic gene expression (both intrinsic and extrinsic noise) can be modeled using SDEs with distributions of parameter values reflecting cellular heterogeneity. To address heterogeneities in gene dynamics across different cells, we propose an END-Nsde method that is able to reconstruct a family of SDEs for the same gene expression process under different parameters. Specifically, for a given set of (biological) parameters *ω*, we are interested in reconstructing

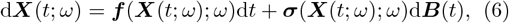

using the approximate SDE

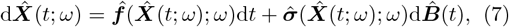

in the sense that the errors 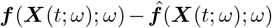 and 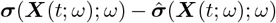 for all different values of *ω* can be minimized. In Eq. (7), 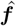 and 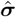 are represented by two parameterized neural networks that take both the state variable 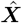 and the parameters *ω* as inputs. To train these two neural networks, we propose an extrinsic-noise-driven temporally decoupled squared *W*_2_ distance loss function

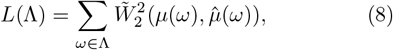

where *µ*(*ω*) and 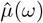 are the distributions of the trajectories ***X***(*t*; *ω*), 0 ≤ *t* ≤ *T* and 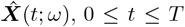,and 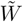 is the temporally decoupled squared *W*_2_ loss function in Eq. (3). Λ denotes the set of parameters *ω*. Note that Eq. (8) is different from the local squared *W*_2_ loss in Refs. [34, 35] since we do not require a continuous dependence of {***X***(*t*; *ω*)}_*t*∈ [0,*T*]_ on the parameter *ω* nor do we require that *ω* is a continuous variable. The extrinsic-noise-driven temporally decoupled squared *W*_2_ loss function Eq. (8) takes into account both parameter heterogeneity and intrinsic fluctuations as a result of the Wiener process ***B***(*t*) and 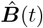 in Eqs. (1) and (2).

We summarize the END-nSDE method in Figs. 1A, B. With observed noisy single-cell dynamic trajectories as the training data, we train two parameterized neural networks by minimizing Eq. (8) to approximate the drift and diffusion terms in the SDE. The reconstructed nSDE is a surrogate model of single-cell dynamics (see Figs. 1 A, B). The hyperparameters and settings for training the neural SDE model are summarized in Table II of Appendix A.

### B. Biological models

We consider three biological examples where stochastic dynamics play a critical role and use our END-nSDE method to reconstruct noisy single-cell gene expression dynamics under both intrinsic and extrinsic noise (also summarized in Fig. 1C). In these applications, we investigate the extent to which the END-nSDE can efficiently capture and infer changes in the dynamics driven by extrinsic noise.

#### 1. Noisy oscillatory circadian clock model

Circadian clocks, often with a typical period of approximately 24 hours, are ubiquitous in intrinsically noisy biological rhythms generated at the single-cell molecular level [36].

We consider a minimal SDE model of the periodic gene dynamics responsible for *per* gene expression which is critical in the circadian cycle. Since *per* gene expression is subject to intrinsic noise [37], we describe it using a linear damped-oscillator SDE

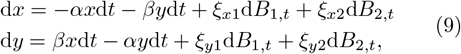

where *x* and *y* are the dimensionless concentrations of the *per* mRNA transcript and the corresponding *per* protein, respectively. d*B*_1,*t*_, d*B*_2,*t*_ are two independent Wiener processes and the parameters *α >* 0 and *β >* 0 denote the damping rate and angular frequency, respectively. A stability analysis at the steady state (*x, y*) = (0, 0) in the noise-free case (*ξ*_*x*_ = *ξ*_*y*_ = 0 in Eq. (9)) reveals that the real parts of the eigenvalues of the Jacobian matrix 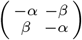 at (*x, y*) = (0, 0) are all negative, indicating that the origin is a stable steady state when the system is noise-free. Noise prevents the state (*x*(*t*), *y*(*t*)) from staying at (0, 0); thus, fluctuations in the single-cell circadian rhythm is noise-induced [37].

To showcase the effectiveness of our proposed END-nSDE method, we take different forms of the diffusion functions *ξ*_*x*_ and *ξ*_*y*_ in Eq. (9), accompanied by different values of noise strength and the correlation between the diffusion functions in the dynamics of *x, y*.

#### 2. RPA-DNA binding model

Regulation of gene expression relies on complex interactions between proteins and DNA, often described by the kinetics of binding and dissociation. Replication protein A (RPA) plays a pivotal role in various DNA metabolic pathways, including DNA replication and repair, through its dynamic binding with single-stranded DNA (ssDNA) [38–41]. By modulating the accessibility of ssDNA, RPA regulates multiple biological mechanisms and functions, acting as a critical regulator within the cell [42]. Understanding the dynamics of RPA-ssDNA binding is therefore a research area of considerable biological interest and significance.

Multiple binding modes and volume exclusion effects complicate the modeling of RPA-ssDNA dynamics. The RPA first binds to ssDNA in 20 nucleotide (nt) mode, which occupies 20nt of the ssDNA. When the subsequent 10nt of ssDNA is free, 20nt-mode RPA can transform to 30nt-mode, further stabilizing its binding to ssDNA, as illustrated in Fig. 2. Occupied ssDNA is not available for other proteins to bind. Consequently, the gap size between adjacent ssDNA-bound RPAs determines the ss-DNA accessibility to other proteins.

**Fig 2.**
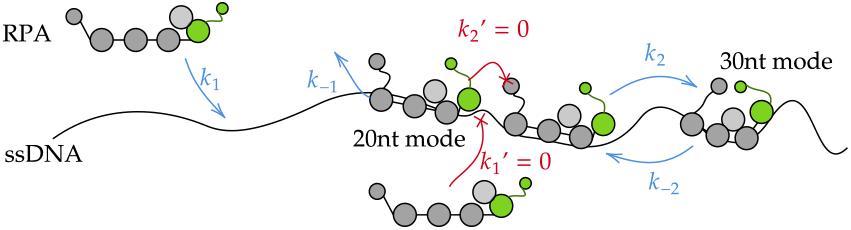
A continuous-time discrete Markov chain model for multiple RPA molecules binding to long ssDNA. The possible steps in the biomolecular kinetics of RPA on ssDNA create a complex scenario involving multiple RPA molecules binding. The RPA in the free solution can bind to ssDNA with rate *k*_1_ provided there are at least 20 nucleotides (nt) of consecutive unoccupied sites. This bound “20-nt mode” RPA unbinds with rate *k*_−1_. When space permits, the 20nt-mode RPA can extend and bind an additional 10nt of DNA at a rate of *k*_2_, converting it to a 30nt mode bound protein. The 30nt-mode RPA transforms back to 20nt-mode spontaneously with the rate *k*_−2_. However, when the gap is not large enough to accommodate the RPA, the binding or conversion is prohibited (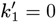 and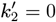).

Mean-field mass-action type chemical kinetic ODE models cannot describe the process very well because they do not capture the intrinsic stochasticity. A stochastic model that tracks the fraction of two different binding modes of RPA, 20nt-mode (*x*_1_) and 30nt-mode (*x*_2_), has been developed to capture the dynamics of this process. A brute-force approach using forward stochastic simulation algorithms (SSAs) [43] was then used to fit the model to experimental data [42]. However, a key challenge in this approach is that the model is nondifferentiable with respect to the kinetic parameters, making it difficult to estimate parameters. Yet, simple spatially homogeneous stochastic chemical reaction systems can be well approximated by a corresponding SDE of the form given in Eq. (1) when the variables are properly scaled in the large system size limit [44]. While interparticle interactions shown in Fig. 2 make it difficult to find a closed-form SDE approximation, the results in Ref. [44] motivate the possibility of an SDE approximation for the RPA-ssDNA binding model in terms of the variables *x*_1_ and *x*_2_.

Here, to address the non-differentiability issue associated with the underlying Markov process, we use our END-nSDE model to construct a differentiable surrogate for SSAs, allowing it to be readily trained from data. Further details on the models and data used in this study are provided in Appendix B. Throughout our analysis of RPA-DNA binding dynamics, we benchmark the SDE reconstructed by our extended *W*_2_-distance approach against those found using other time series analysis and reconstruction methods such as the Gaussian process, RNN, LSTM, and the neural ODE model. We show that our surrogate SDE model is most suitable for approximating the RPA-DNA binding process because it can capture the intrinsic stochasticity in the dynamics.

### 3. NFκB signaling model

Macrophages can sense environmental information and respond accordingly with stimulus-response specificity encoded in signaling pathways and decoded by downstream gene expression profiles [45]. The temporal dynamics of NF*κ*B, a key transcription factor in immune response and inflammation, encodes stimulus information [46]. NF*κ*B targets and regulates vast immune-related genes [47–49]. While NF*κ*B signaling dynamics are stimulus-specific, they exhibit significant heterogeneity across individual cells under identical conditions [46]. Understanding how specific cellular heterogeneity (extrinsic noise) contributes to heterogeneity in NF*κ*B signaling dynamics can provide insight into how noise affects the fidelity of signal transduction in immune cells. A previous modeling approach employs a 52-dimensional ODE system to quantify the NF*κ*B signaling network [46] and recapitulate the signaling dynamics of a representative cell. This ODE model includes 52 molecular entities and 47 reactions across a TNF-receptor module, an adaptor module, and a core module with and NF*κ*B-IKK-I*κ*B*α* (I*κ*B*α* is an inhibitor of NF*κ*B, while IKK is the I*κ*B kinase complex that regulates the I*κ*B*α* degradation) feedback loop (see Fig. 3) [50]. However, such an ODE model is deterministic and assumes no intrinsic fluctuations in the biomolecular processes. Yet, from experimental data, the NF*κ*B signaling dynamics fluctuate strongly; such fluctuations cannot be quantitatively described by any deterministic ODE model. Due to the system’s high dimensionality and nonlinearity, it is challenging to quantify how intrinsic noise influences temporal coding in NF*κ*B dynamics.

**Fig 3.**
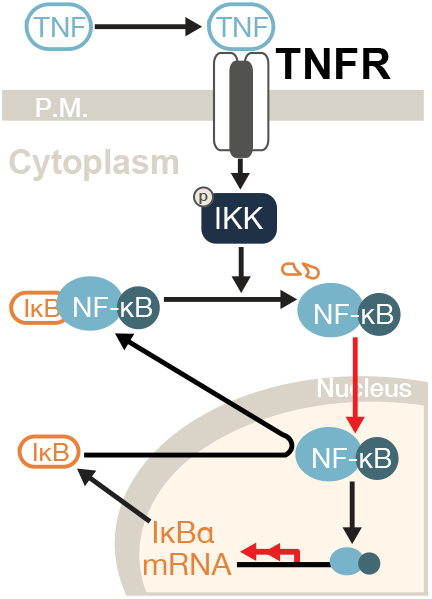
Simplified schematic of the NF*κ*B Signaling Network. TNF binds its receptor, activating IKK, which degrades I*κ*B*α* and releases NF*κ*B. The free NF*κ*B translocates to the nucleus and promotes I*κ*B*α* transcription. Newly synthesized I*κ*B*α* then binds NF*κ*B and exports it back to the cytoplasm. Red arrows indicate noise that we consider in the corresponding SDE system.

To incorporate the intrinsic noise within the NF*κ*B signaling network, we introduce noise terms into the 52-dimensional ODE system to build an SDE that can account for the observed temporally fluctuating nuclear NF*κ*B trajectories. While NF*κ*B signaling pathways involve many variables, experimental constraints limit the number of measurable components. Among these, nuclear NF*κ*B activity is the most direct and critical experimental readout. As a minimal stochastic model, we hypothesize that only the biophysical and biochemical processes of NF*κ*B translocation (which directly affects experimental measurements) and I*κ*B*α* transcription (a key regulator of NF*κ*B translocation) are subject to Brownian-type noise (red arrows in Fig. 3), as these processes play crucial roles in the oscillatory dynamics of NF*κ*B [50].

The intensity of Brownian-type noise in the NF*κ*B dynamics may depend on factors such as cell volume (smaller volumes result in higher noise intensity), or copy number (lower copy numbers lead to greater noise intensity), and is therefore considered a form of extrinsic noise. Noise intensity parameters thus capture an aspect of cellular heterogeneity. There are other sources of cellular heterogeneity, such as variations in kinase or enzyme abundances, which are too complicated to model and are thus not included in the current model. For simplicity, all kinetic parameters, except for the noise intensity (*σ*), are assumed to be consistent with those of a representative cell [50]. The 52-dimensional ODE model for describing NF*κ*B dynamics is given in Refs. [46, 51]. We extend this model by adding noise to the dynamics of the sixth, ninth, and tenth ODEs of the 52-dimensional ODE model. We retain 49 ODEs but convert the equations for the sixth, ninth, and tenth components to SDEs:

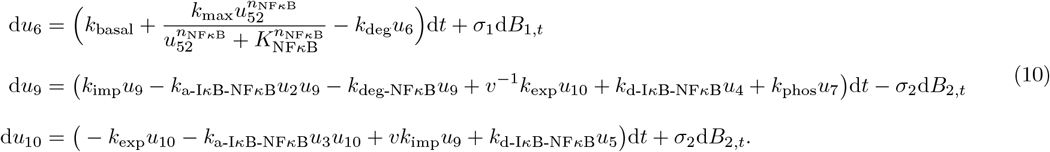

In Eqs. 10, *u*_2_ is the concentration of I*κ*B*α*; *u*_3_ is the concentration of I*κ*B*α*; *u*_4_ is the concentration of the I*κ*B*α*-NF*κ*B complex; *u*_5_ is the concentration of the I*κ*B*α*-NF*κ*B complex in the nucleus; *u*_6_ is the mRNA of I*κ*B*α*; *u*_7_ is the IKK-I*κ*B*α*-NF*κ*B complex; *u*_9_ is NF*κ*B; *u*_10_ represents nuclear NF*κ*B activity; and *u*_52_ is the nuclear concentration of NF*κ*B with RNA polymerase II that is ready to initiate mRNA transcription. A description of the parameters and their typical values are given in the supplemental table IV. The quantities *σ*_1_d*B*_1,*t*_ and *σ*_2_d*B*_2,*t*_ are noise terms associated with I*κ*B*α* transcription and NF*κ*B translocation, respectively. The remaining variables are latent variables and their dynamics are regulated via the remaining 49-dimensional ODE in Refs. [46, 51]. The activation of NF*κ*B is quantified by the nuclear NF*κ*B concentration (*u*_5_ + *u*_10_), which is also measured in experiments.

Within this example, we wish to determine if our proposed parameter-associated nSDE can accurately reconstruct the dynamics underlying experimentally observed NF*κ*B trajectory data.

## III. RESULTS

### A. Accurate reconstruction of circadian clock dynamics

As an illustrative example, we use the *W*_2_-distance nSDE reconstruction method to first reconstruct the minimal model for damped oscillatory circadian dynamics (see Eq. (9)) under different forms of the diffusion function. We set the two parameters *α* = 0.19 and *β* = 0.21 in Eq. (9) and impose three different forms for the diffusion functions *ξ*_*x*1_, *ξ*_*x*2_, *ξ*_*y*1_, *ξ*_*y*2_: a constant diffusion function [52], a Langevin [53] diffusion function, and a linear diffusion function [54]. These functions, often used to describe fluctuating biophysical processes, are

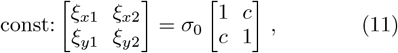

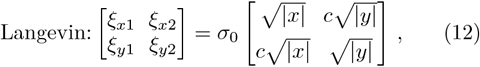

and

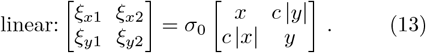

There are two additional parameters in Eqs. (11), (12), and (13): *σ*_0_ that determines the intensity of the Brownian-type fluctuations and *c* that controls the correlation of fluctuations between the two dimensions. For each type of diffusion function, we trained a different nSDE model, each of which takes the state variables (*x, y*) and the two parameters (*c, σ*_0_) as inputs and which outputs the values of the reconstructed drift and diffusion functions.

We take 25 combinations of (*σ*_0_, *c*) ∈{(0.1+0.05*i*, 0.2+ 0.2.2*j*), *i* ∈ *{*0, …, 4*}, j* ∈ *{*0, …, 4*}*; for each combination of (*ξ*_*x*1_, *ξ*_*x*2_, *ξ*_*y*1_, *ξ*_*y*2_), we generate 50 trajectories from the ground truth SDE (9) as the training data with *t*∈ [0, 1]. The initial condition is set as (*x*(0), *y*(0)) = (0, 1). To test the accuracy of the reconstructed diffusion and drift functions, we measure the following relative errors:

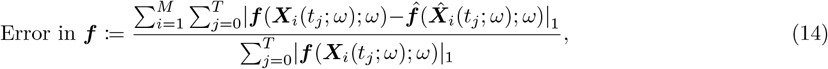

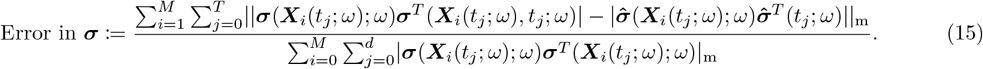

Here, ***f*** := (−*αx* −*βy, βx*− *αy*)^*T*^ is the vector of ground truth drift functions and 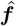 is the reconstructed drift function. ***σ*** is the matrix of ground truth diffusion functions [*ξ*_*x*1_, *ξ*_*x*2_; *ξ*_*y*1_, *ξ*_*y*2_] given in Eqs. (11), (12), and (13). *M* is the number of training samples, | *·* |_1_ denotes the ℓ^1^ norm of a vector, and the matrix norm 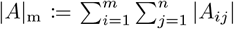 for a matrix *A* ∈ ℝ^*m×n*^. The errors are measured separately for different parameters *ω* := (*σ*_0_, *c*).

The errors in the reconstructed drift function 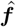 and diffusion function 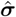 as well as the temporally decoupled squared *W*_2_ loss Eq. (3) associated with different forms of the diffusion function and different values of (*σ*_0_, *c*) are shown in Fig. 4. When the diffusion function is a constant Eq. (11), the mean reconstruction error of the drift function is 0.15, the mean reconstruction error of the diffusion function is 0.16, and the mean temporally decoupled squared *W*_2_ loss between the ground truth trajectories and the predicted trajectories is 0.074 (averaged over all sets of parameters (*σ*_0_, *c*)). When a Langevin-type diffusion function Eq. (12) is used as the ground truth, the mean errors for the reconstructed drift and diffusion functions are 0.069 and 0.29, respectively, and the mean temporally decoupled squared *W*_2_ loss between the ground truth and predicted trajectories is 0.020. For a linear-type diffusion function as the ground truth, mean reconstruction errors of the drift and diffusion functions are 0.19 and 0.41, respectively, and the mean temporally decoupled squared *W*_2_ distance is 0.013. For all three forms of diffusion, our END-nSDE method can accurately reconstruct the drift function (−*αx* −*βy, βx*− *αy*) (see Figs. 4D-F). When the diffusion function is a constant, our END-nSDE model can also accurately reconstruct this constant (see Fig. 4G). When the diffusion function takes a more complicated form such as the Langevin-type diffusion function Eq. (12) or the linear-type diffusion function Eq. (13), the reconstructed nSDE model can still approximate the diffusion function well for most combinations of (*c, σ*_0_), especially when the correlation *c >* 0.2 (see Figs. 4H-I). Overall, our proposed END-nSDE model can accurately reconstruct the minimal stochastic circadian dynamical model Eq. (9) in the presence of extrinsic noise (different values of (*σ*_0_, *c*)); the accuracy of the reconstructed drift and diffusion functions is maintained for most combinations of (*σ*_0_, *c*).

**Fig 4.**
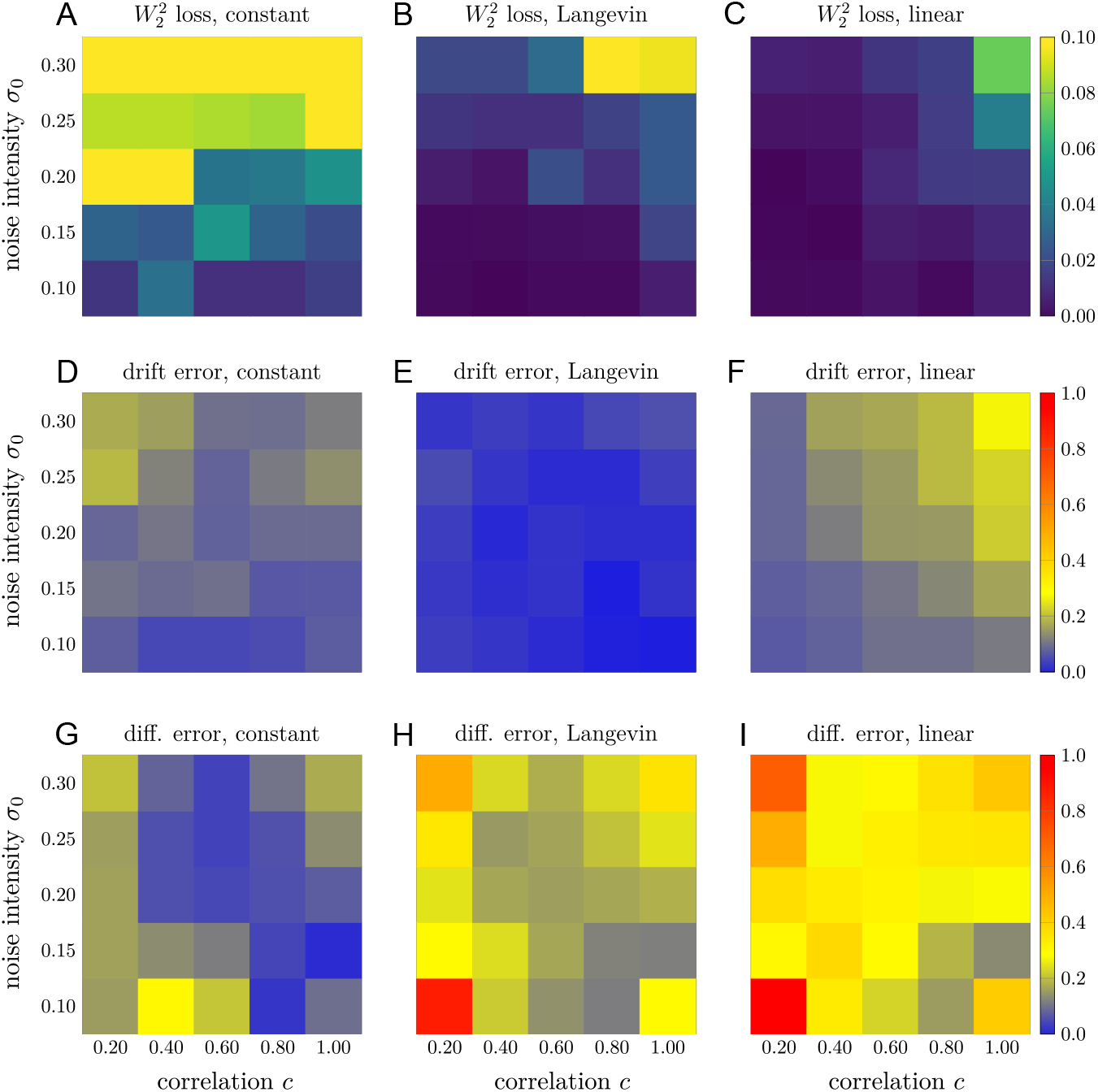
Reconstructing the circadian model using END-nSDE. Temporally decoupled squared *W*_2_ losses Eq. (3) and errors in the reconstructed drift and diffusion functions for different types of the diffusion function and different values of (*σ*_0_, *c*). A-C. The temporally decoupled squared *W*_2_ loss between the ground truth trajectories and the trajectories generated by the reconstructed nSDEs for the constant-type diffusion function Eq. (11), Langevin-type diffusion function Eq. (12), and the linear-type diffusion function Eq. (13). D-F. Errors in the reconstructed drift function for the three different types of ground truth diffusion functions and the linear-type diffusion function Eq. (13). G-I. Errors in the reconstructed diffusion function for the three different types of ground truth diffusion functions.

### B. Accurate approximation of interacting DNA-protein systems with different kinetic parameters

To construct a differentiable surrogate for stochastic simulation algorithms (SSAs), the neural SDE model should be able to take kinetic parameters as additional inputs. Thus, the original *W*_2_-distance SDE reconstruction method in [29] can no longer be applied because the trained neural SDE model cannot take into account extrinsic noise, *i*.*e*., different values of kinetic parameters. To be specific, we vary one parameter (the conversion rate *k*_2_ from 20nt-mode RPA to 30nt-mode RPA) in the stochastic model and then apply our END-nSDE method which takes the state variables and the kinetic parameter *k*_2_ as the input. We set *k*_2_ ∈ {10^−4+*j/*10^, *j* = 0, …, 25} with other parameters taken from experiments [42] (*k*_−1_ = 10^−3^ s^−1^, *k* _− 1_ = 10^−6^ s^−1^, *k*_− 2_ = 10^−6^ s^−1^, see Fig. 2). For each *k*_2_, we generate 100 trajectories and use 50 for the training set and the other 50 for the testing set. Each trajectory encodes the dynamics of the fraction of 20-nt mode DNA-bound RPA *x*_1_(*t*) and the fraction of 30-nt mode DNA-bound RPA *x*_2_(*t*).

When approximating the dynamics underlying the RPA-DNA binding process, we compare our SDE reconstruction method with other benchmark time-series analysis or reconstruction approaches, including the RNN, LSTM, Gaussian process, and the neural ODE model. These benchmarks are described in detail in Appendix C 2.

The extrinsic-noise-driven temporally decoupled squared *W*_2_ distance loss Eq. (8) between the distribution of the ground truth trajectories and the distribution of the predicted trajectories generated by our END-nSDE reconstructed SDE model is the smallest among all methods (shown in Table I). The underlying reason is an SDE well approximates the genuine Markov counting process process underlying the continuum-limit RPA-DNA binding process [44]. The RNN and LSTM models do not capture the intrinsic fluctuations in the counting process. The neural ODE model is a deterministic model and cannot capture the stochasticity in the RPA-DNA binding dynamics. Additionally, the Gaussian process can only accurately approximate linear SDEs, which is not an appropriate form for an SDE describing the RPA-DNA binding process.

**TABLE I.** The extrinsic-noise-driven time-decoupled squared *W*_2_ distance Eq. (8) between the ground truth and predicted trajectories generated by different models on the testing set.

| Model | Loss |
| --- | --- |
| END-nSDE | **0.0006** |
| LSTM | 0.062 |
| RNN | 0.087 |
| nODE | 0.0012 |
| Gaussian Process | 0.0010 |

In Figs. 5A,B, we plot the predicted trajectories obtained by the trained neural SDE model for two different values lg *k*_2_ = −4 and lg *k*_2_ = −1.5. Actually, for all different values of *k*_2_, trajectories generated by our END-nSDE method match well with the ground truth trajectories on the testing set, as the temporally decoupled squared *W*_2_ loss is maintained small for all *k*_2_ (shown in Fig. 5C). This demonstrates the ability of our method to capture the dependence of the stochastic dynamics on biochemical kinetic parameters.

**Fig 5.**
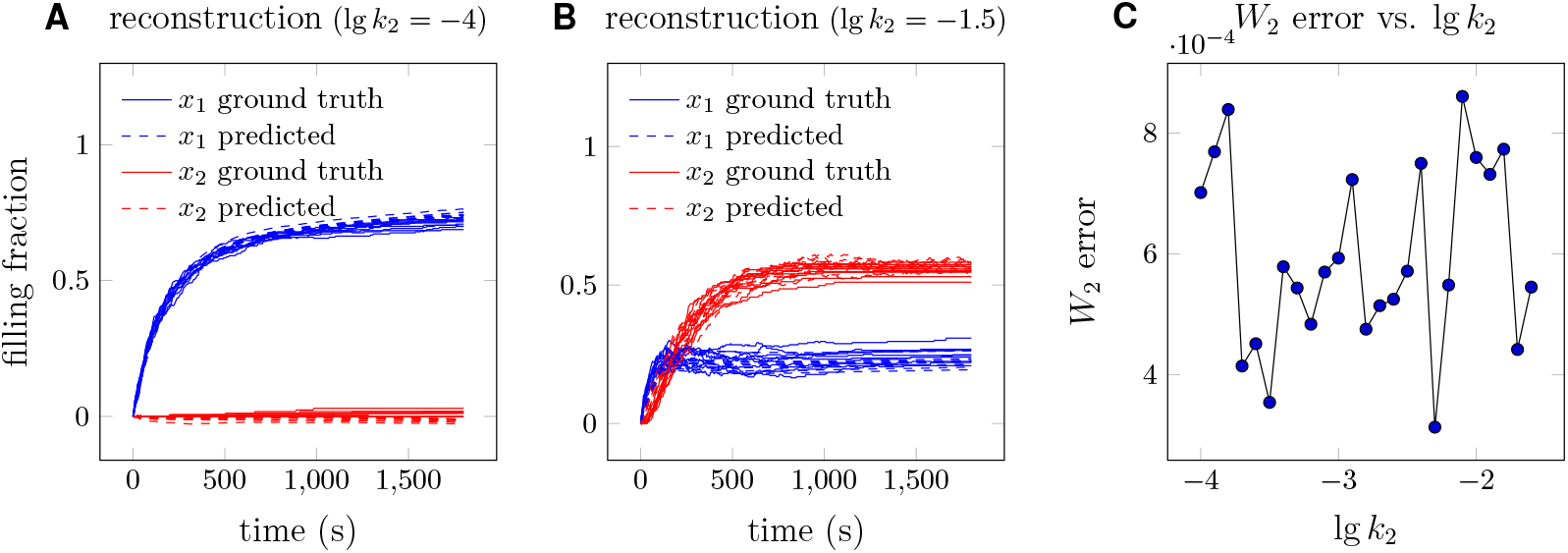
Reconstructed trajectories of the RPA-DNA binding problem. A. Sample ground truth and reconstructed trajectories evaluated at lg *k*_2_ = − 4, where we use the convention that lg = log10. B. Sample ground truth and reconstructed parameters evaluated at lg *k*_2_ = − 1.5. C. Temporally decoupled squared *W*_2_ distances (see Eq. (8)) between the ground truth and reconstructed trajectories evaluated at different lg *k*_2_ values. In A and B, blue and red trajectories represent the filling fractions of DNA by 20nt-mode and 30nt-mode RPA, respectively. The dashed lines represent the predicted trajectories, and the solid lines represent the ground truth. Throughout the figure, the data are generated by a single neural SDE model that accepts the conversion rate *k*_2_ as a parameter and outputs the trajectories.

### C. Reconstructing high-dimensional NF*κ*B signaling dynamics from simulated and experimental data

Finally, we evaluate the effectiveness of the END-nSDE framework in reconstructing high-dimensional NF*κ*B signaling dynamics under varying noise intensities and investigate the performance of the neural SDE method in reconstructing experimentally measured noisy NF*κ*B dynamics. The procedure is divided into two parts. First, we trained and tested our END-nSDE method on synthetic data generated by the NF*κ*B SDE model Eq. (10) under different noise intensities (*σ*_1_, *σ*_2_). Second, we test whether the trained END-nSDE can reproduce the experimental dynamic trajectories.

#### 1. Reconstructing a 52-dimensional stochastic model for NFκB dynamics

For training END-nSDE models, we first generate synthetic data from the 52-dimensional SDE model of NF*κ*B signaling dynamics Eqs. (10) and established models [46, 51]. The synthetic trajectories are generated under 121 combinations of noise intensity (*σ*_1_, *σ*_2_) in Eqs. (10) (see Appendix D). The resulting NF*κ*B trajectories vary depending on noise intensity, with low-intensity noise producing more consistent dynamics across cells (see Fig. 6A) and higher-intensity noise yielding more heterogeneous dynamics (see Fig. 6B). The simulated ground truth trajectories are split into training and testing datasets (see Appendix E for details). Specifically, we exclude 25 combinations of noise intensities (*σ*_1_, *σ*_2_) from the training set in order to test the generalizability of the trained neural SDE model on noisy intensities.

**Fig 6.**
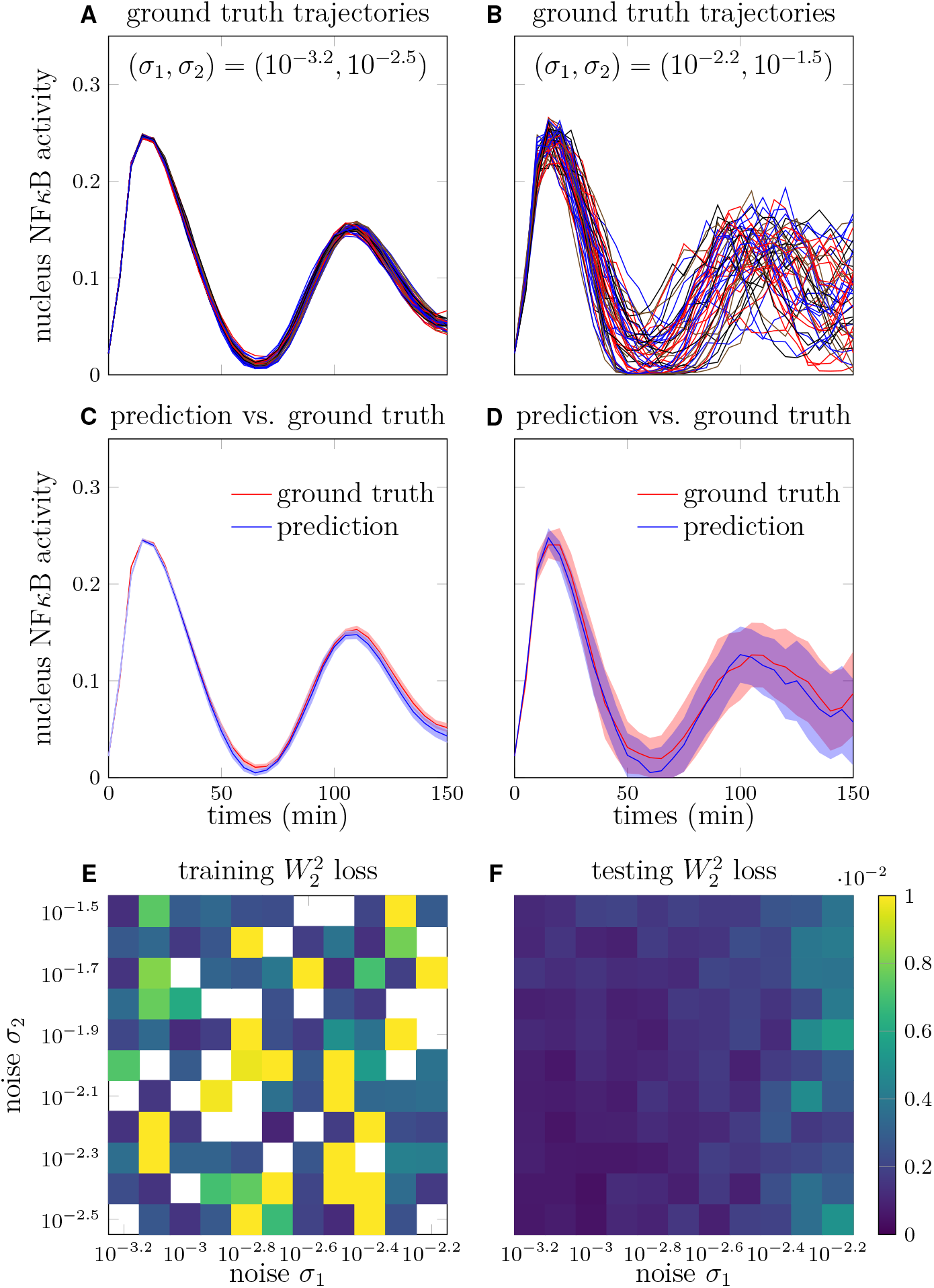
Reconstruction of NF*κ*B signaling dynamics. A. Sample trajectories of nuclear NF*κ*B concentration as a function of time with *σ*_1_ = 10^−3.2^, *σ*_2_ = 10^−2.5^. B. Sample trajectories of nuclear NF*κ*B concentration as a function of time with *σ*_1_ = 10^−2.2^, *σ*_2_ = 10^−1.5^. C. Reconstructed nuclear NF*κ*B trajectories generated by the trained neural SDE versus the ground truth nuclear NF*κ*B trajectories under noise intensities *σ*_1_ = 10^−3.2^, *σ*_2_ = 10^−2.5^ in Eqs. (10). D. Reconstructed nuclear NF*κ*B trajectories generated by the trained neural SDE versus the ground truth nuclear NF*κ*B trajectories under noise intensities *σ*_1_ = 10^−2.2^, *σ*_2_ = 10^−1.5^. E. The squared *W*_2_ distance between the distributions of the predicted trajectories and ground truth trajectories on the training set under different noise strengths (*σ*_0_, *σ*_1_). For training, we randomly selected 50% sample trajectories in 80 combinations of noise strengths (*σ*_1_, *σ*_2_) as the training dataset. Blank cells indicate that the corresponding parameter set is not included in the training set. F. Validation of the trained model by evaluating the squared *W*_2_ distance between the distributions of predicted trajectories and ground truth trajectories on the validation set.

Next, we trained a 52-dimensional neural SDE model using our END-nSDE method on synthetic trajectories (see Appendix E for details). The loss function is based on the *W*_2_ distance between the distributions of the neural SDE predictions in Eqs. (10) and the simulated nuclear NF*κ*B and I*κ*B*α*-NF*κ*B complex activities (*u*_5_(*t*) and *u*_10_(*t*), respectively) and the corresponding END-nSDE predictions. The remaining 50 variables of the NF*κ*B system are treated as latent variables, as they are not directly included in the loss function calculation.

Although the NF*κ*B dynamics vary under different noise intensities (*σ*_1_, *σ*_2_), the trajectories generated by our trained neural SDE closely align with the ground truth synthetic NF*κ*B dynamics under different noise intensities (*σ*_1_, *σ*_2_) (see Figs. 6C-D). The neural SDE model demonstrates greater accuracy in reconstructing NF*κ*B dynamics when the noise in I*κ*B*α* transcription (*σ*_1_) is smaller, as evidenced by the reduced squared *W*_2_ distance between the predicted and ground-truth trajectories on both the training and validation sets (see Figs. 6E-F). The temporally decoupled squared *W*_2_ loss Eq. (8) on the validation set is close to that on the training set for different values of noise intensities (*σ*_1_, *σ*_2_). The mean squared *W*_2_ distance across all combinations of noise intensities (*σ*_1_, *σ*_2_) is 0.0013 for the training set, and the validation set shows a mean squared *W*_2_ distance of 0.0017.

Since the loss function for this application involves only two variables out of 52, we also tested whether the “full” 52-dimensional NF*κ*B system can be effectively modeled by a two-dimensional neural SDE. After training, we found that the reduced model was insufficient for reconstructing the full 52-dimensional dynamics, as it disregarded the 50 latent variables not included in the loss function (see Fig. S4 in Appendix F). This result underscores the importance of incorporating latent variables from the system, even when they are not explicitly included in the loss function.

#### 2. Reconstructing NFκB experimental data with END-nSDE

We then assess whether our proposed END-nSDE can accurately reconstruct the experimentally measured NF*κ*B dynamic trajectories. For simplicity and feasibility, we tested the END-nSDE under the assumption that(1) all cells share the same drift function, and (2) cells with trajectories that deviate similarly from their ODE predictions have the same noise intensities. Based on these assumptions, we developed the following workflow (see Fig. 7):

**Fig 7.**
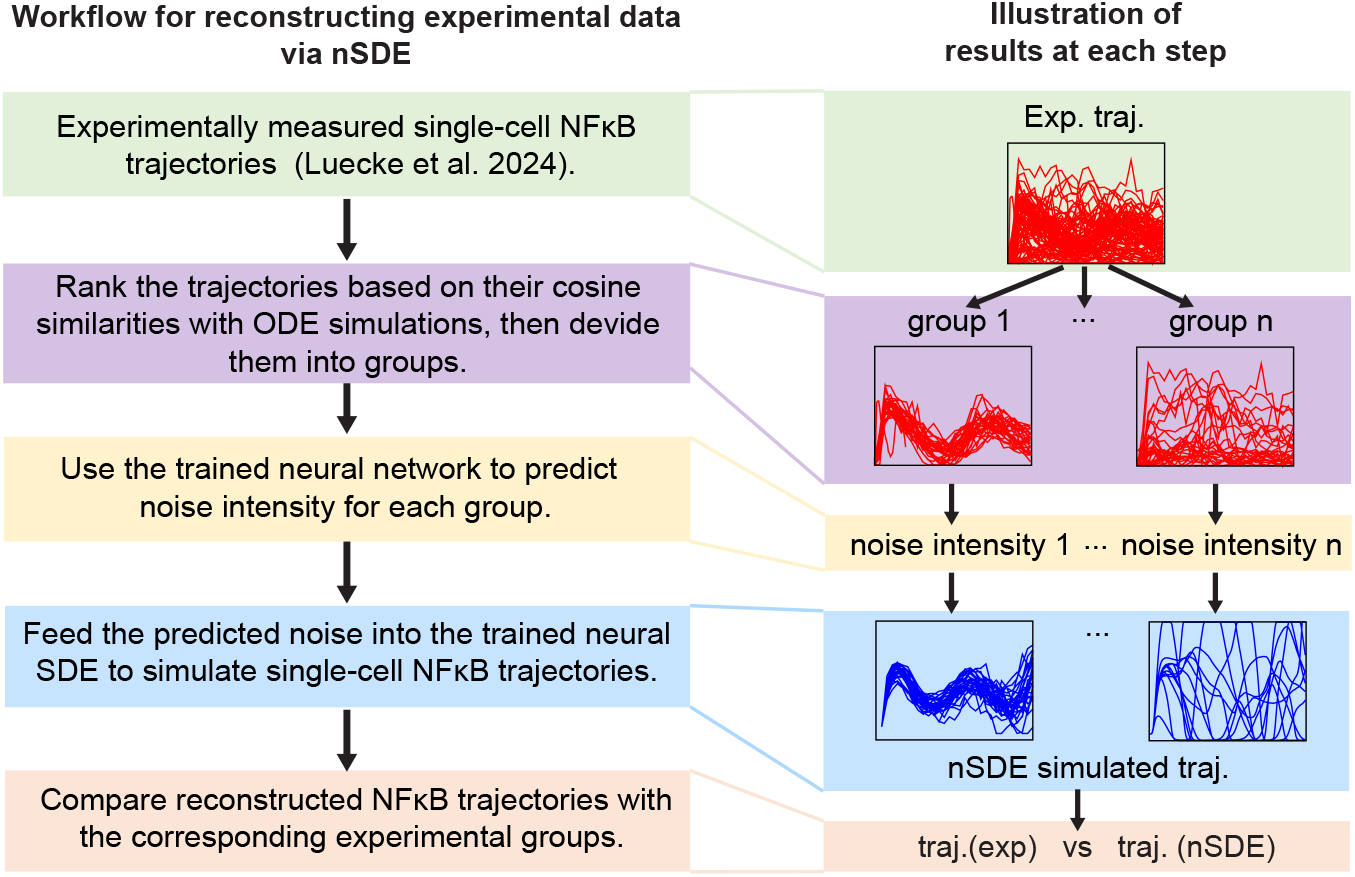
Workflow of reconstructing experimental data via END-nSDE. Workflow for reconstructing experimental data using the trained parameterized nSDE and the parameter-inference neural network (NN). The boxes on the left outline the steps of the experimental data reconstruction process, while the boxes on the right illustrate the corresponding results at each step.

1. We used experimentally measured single-cell trajectories of NF*κ*B activity, obtained through live-cell image tracking of macrophages from mVenus-tagged RelA mouse with a frame frequency of five minutes [55]. These trajectories correspond to the sum of nuclear I*κ*B*α* − NF*κ*B and NF*κ*B in the 52D SDE model (*u*_5_(*t*) and *u*_10_(*t*) in Eqs. (10)).
2. The experimental dataset was divided into subgroups. Cosine similarity was calculated between the ODE-generated trajectory (representative-cell NF*κ*B dynamics) and experimental trajectories. The trajectories were then ranked and divided into different groups based on their cosine similarity with the ODE model. Experimental trajectories with higher similarity to the ODE trajectory are expected to exhibit smaller intrinsic fluctuations, corresponding to lower noise intensities (see Appendix G for details).
3. Each group of experimental trajectories was input into the trained neural network (see the next paragraph for more details) to infer the corresponding noise intensities (*σ*_1_, *σ*_2_). For simplicity, we assume that trajectories within each group shared the same noise intensities.
4. The inferred noise is then used as inputs for the trained END-nSDE to simulate NF*κ*B trajectories.
5. The simulated trajectories were compared with the corresponding experimental data to evaluate the model’s performance.

To estimate noise intensities from different groups of experimentally measured single-cell nuclear NF*κ*B trajectories (step (3) in the proposed workflow), we trained another neural network to predict the corresponding I*κ*B*α* transcription and NF*κ*B translocation noise intensities from the groups of NF*κ*B trajectories in the synthetic training data. The trained neural network can then be used for predicting noise intensities in the validation set (see Appendix H for details).

Assessing the impact of group size (number of trajectories) on noise intensity prediction performance, we found that taking a group size of at least two leads to a relative error of around 0.1 (see Fig. 8A). Given the high heterogeneity present in experimental data, we took a group size of 32 as the input into the neural network. Under this group size, the relative errors in the predicted noise intensities were 0.021 on the training set and 0.062 on the testing set (see Figs. 8B-C).

**Fig 8.**
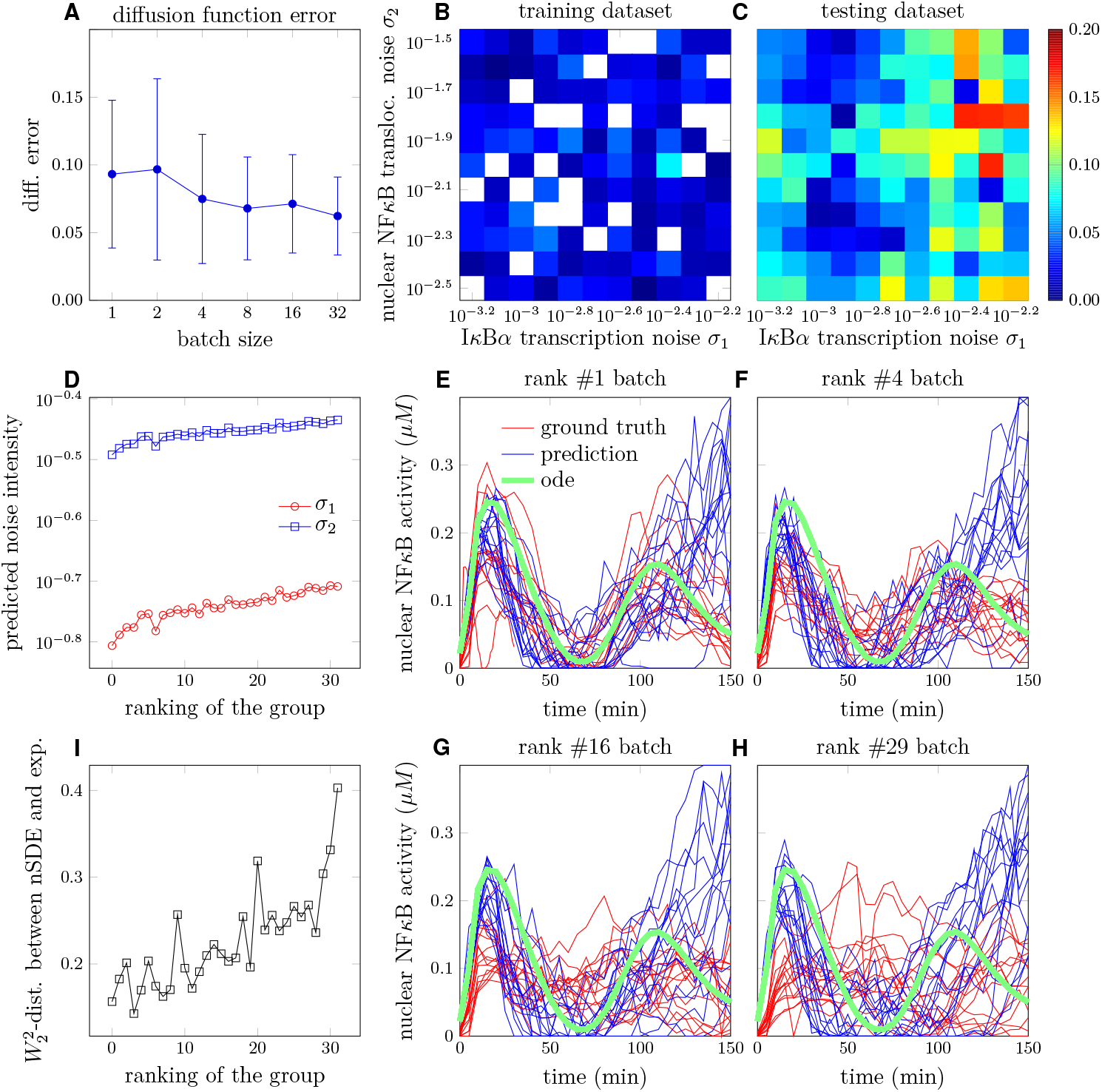
Inferring intrinsic noise intensities and reconstructing experimental data via END-nSDE. A. Plots showing the mean (solid circles) and variance (error bars) of the relative error in the reconstructed noise intensities 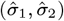 predicted by the parameter-inference NN for the testing dataset, as a function of the group size of input trajectories. B. Heatmaps showing the relative error in the reconstructed noise intensities for the training dataset. Colored cells represent results from the parameter-inference NN for the training dataset, while blank cells indicate noise strength values not included in the training set. C. Heatmaps showing the relative error in the diffusion function for the testing dataset. D. The inferred intensity of I*κ*B*α* transcription noise (*σ*_1_) and NF*κ*B translocation noise (*σ*_2_) in different groups of experimental trajectories, plotted against the group’s ranking in decreasing similarity with the representative ODE trajectory. E-H. Groups of experimental and nSDE-reconstructed trajectories ranked by decreasing cosine similarity: #1 (E), #4 (F), #16 (G), #29 (H). The squared *W*_2_-distance between experimental and SDE-generated trajectories are 0.157 (E), 0.143 (F), 0.212 (G), 0.236 (H). The inferred noises are (10^−0.49^, 10^−0.81^) (E), (10^−0.47^, 10^−0.78^) (F), (10^−0.46^, 10^−0.74^) (G), (10^−0.44^, 10^−0.71^) (H). I. The temporally decoupled squared *W*_2_ distance between reconstructed trajectories generated by the trained END-nSDE and groups of experimental trajectories, ordered according to decreasing similarity with the representative ODE trajectory.

Using the trained neural network, we inferred noise intensities for the experimental data, which were grouped based on their cosine similarities with the representative-cell trajectory (deterministic ODE) with a group size of 32. The predicted noise intensities on the experiment data set are larger than the noise intensities on the training set, and the underlying reason could be extrinsic noise which is not taken into account interferes with the inference of noise intensity. The transcription noise of I*κ*B*α* is predicted to be within the range of [10^−0.81^, 10^−0.71^] (see Fig. 8D). In addition, the inferred noise for NF*κ*B translocation fell within [10^−0.49^, 10^−0.43^] (see Fig. 8D). These inferred noise intensities were then used as inputs to the END-nSDE to simulate NF*κ*B trajectories.

We compare the reconstructed NF*κ*B trajectories generated by the trained neural SDE model with the experimentally measured NF*κ*B trajectories (see Figs. 8E-I). The trajectories generated using our END-nSDE method successfully reproduce the experimental dynamics for the majority of time points for the top 50% of cell subgroups most correlated with the representative-cell ODE model (see Figs. 8E-G, Figs. 8I).

For the top-ranked subgroups (#1 to #16), the heterogeneous nSDE-reconstructed dynamics align well with the experimental data for the first 100 minutes. The predicted trajectories deviate more from ground truth trajectories observed in experiments after 100 minutes possibly due to error accumulation and errors in the predicted noise intensity. For experimental subgroups that significantly deviate from the representative-cell ODE model, the END-nSDE struggles to fully capture the heterogeneous trajectories. This limitation likely arises from the assumption that all cells in a group share the same underlying dynamics, whereas in reality, substantial cellular differences in underlying dynamics exist due to heterogeneity in the drift term, an aspect not accounted for in END-nSDE due to the high computational cost.

Nonetheless, our END-nSDE can partially reconstruct experimental datasets and has the potential to fully capture experimental dynamics. With sufficient computational resources, our proposed workflow can also incorporate extrinsic noise in the cellular dynamics drift terms, allowing for further discrimination of experimental trajectories.

Overall, we have demonstrated that reconstructing noisy experimental trajectories can be accomplished by (i) inferring noise intensities from noisy trajectories grouped by different noise levels and (ii) using END-nSDE to reconstruct the experimental data based on the inferred noise.

## IV. DISCUSSION

In this work, we used a *W*_2_-distance to develop an END-nSDE reconstruction method that takes into account extrinsic noise in gene expression dynamic as observed across various biophysical and biochemical processes such as circadian rhythms, RPA-DNA binding, and NF*κ*B translocation. We first demonstrated that our END-nSDE method can successfully reconstruct a minimal noise-driven fluctuating SDE characterizing the circadian rhythm, showcasing its effectiveness in reconstructing SDE models that contain both intrinsic and extrinsic noise. Next, we used our END-nSDE method to learn a surrogate extrinsic-noise-driven neural SDE, which approximates the RPA-DNA binding process. Molecular binding processes are usually modeled by a Markov counting process and simulated using Monte-Carlo-type stochastic simulation algorithms (SSAs) [43]. Our END-nSDE reconstruction approach can effectively reconstruct the stochastic dynamics of the RPA-ssDNA binding process while also taking into account extrinsic noise (heterogeneity in biological parameters among different cells). Our END-nSDE method outperforms several benchmark methods such as LSTMs, RNNs, neural ODEs, and Gaussian processes.

Finally, we applied our methodology to analyze NF*κ*B trajectories collected from over a thousand cells. Not only did the neural SDE model trained on the synthetic dataset perform well on the validation set, but it also partially recapitulated experimental trajectories of NF*κ*B abundances, particularly for subgroups with dynamics similar to the representative cell. These results underscore the potential of neural SDEs in modeling and understanding the role of intrinsic noise in complex cellular signaling systems [56– When the experimental trajectories were divided into subgroups, we assumed that all cells across different groups shared the same drift function (as in the representative ODE) and cells within each group shared the same diffusion term. We found that subgroups with dynamics more closely aligned with the deterministic ODE model resulted in better reconstructions. In contrast, for experimental trajectories that deviated significantly from the representative ODE model, their underlying dynamics may differ from those defined by the representative cell’s ODE. Therefore, the assumption that a group shares the same drift function as the representative cell ODE holds only when the trajectories closely resemble the ODE. Incorporating noise into the drift term for training the neural SDE could potentially address this issue. We did not consider this approach due to the high computational cost required for training.

Applying our method to high-dimensional synthetic NF*κ*B datasets, we showed the importance of incorporating latent variables. This necessity arises because the ground-truth dynamics of the measured quantities (nuclear NF*κ*B) are not self-closed and inherently depend on additional variables. Consequently, the 52-dimensional SDE reconstruction requires more variables than just the “observed” dynamics of nuclear NF*κ*B. In this example, the remaining 50 variables in the nSDE were treated as latent variables, even though they were not explicitly included in the loss function.

There are several promising directions for future research. First, it is desirable to identify methods to extract an explicit and interpretable version of the learned neural network SDEs. One might employ a polynomial model for the drift and diffusion functions in the SDE similar to the one suggested in Ref. [59]. Such an explicit representation may facilitate linking neural reconstruction methods to established mechanistic models. The importance of latent variables should also be analyzed to understand if and how they affect the reconstruction of high-dimensional models. The *W*_2_-distance loss landscapes in the examples examined in this study should also be examined in more depth. Prior research [60, 61] has highlighted the importance of studying loss landscapes to characterize and potentially enhance the generalization capabilities of neural networks across various tasks. Finally, neural SDEs can serve as surrogate models for complex biomedical dynamics [62, 63]. Combining such surrogate models with neural control functions [60, 64, 65] can be useful for tackling complex biomedical control problems.

## ACKNOWLEDGMENTS

LB acknowledges financial support from hessian. AI and the ARO through grant W911NF-23-1-0129. TC acknowledges inspiring discussions at the “Statistical Physics and Adaptive Immunity” program the Aspen Center for Physics, which is supported by National Science Foundation grant PHY-2210452. We acknowledge Stefanie Luecke providing the experimental datasets for NF*κ*B dynamics.

## Appendix A: Hyperparameters and conditions used in the neural SDE model

In this section, we provide the hyperparameters in the neural SDE models as well as the training details in Table II.

**TABLE II.** The hyperparameters for training the neural SDE model of each example.

|  | Example 1 | Example 2 | Example 3 |
| --- | --- | --- | --- |
| Gradient descent method | Adam | Adam | Adam |
| Learning rate | 0.002 | 0.002 | 0.002 |
| Weight decay | 0.005 | 0.005 | 0.005 |
| Activation function | ReLU | ReLU | ReLU |
| # of epochs | 500 | 2000 | 2000 |
| # of hidden layers in $\hat{f}$ | 3 | 2 | 2 |
| # of neurons in hidden layer in $\hat{f}$ | 100 | 200 | 400 |
| # of hidden layers in $\hat{\sigma}$ | 3 | 2 | 2 |
| # of neurons in each hidden layer in $\hat{\sigma}$ | 100 | 200 | 400 |
| $\Delta t$ | 0.1 (s) | $\frac{1}{720}$ (hr) | $\frac{1}{12}$ (hr) |
| $T$ | 11 | 361 | 31 |

## Appendix B: RPA dynamic binding to long ssDNA can be simulated by a generalized random sequential adsorption (RSA) model

To elucidate the biophysical mechanism of replication protein A (RPA) binding dynamics to long single-stranded DNA (ssDNA), we developed a continuous-time discrete Markov chain model (see Fig. 2). The Random Sequential Adsorption (RSA) model in a one-dimensional (1D) finite-length context effectively represents the process of protein binding to DNA, capturing the key property that each nucleotide (nt) of ssDNA cannot be occupied by more than one protein molecule. This unique characteristic leads to incomplete occupation even with protein oversaturation, distinguishing DNA-relevant reactions from those described by the mass action law. To reveal finer structures such as gap distribution, we implemented an exact stochastic sampling approach.

We adapted the model based on current knowledge of RPA binding modes, incorporating multiple binding modes and volume exclusion effects. RPA has two binding modes: a 20-nt mode (partial binding mode, PBM) and a 30-nt mode (full-length binding mode, FLBM) (see Fig. 2). RPA initially binds to a 20-nt ssDNA with a rate of *k*_1_ and dissociates at a rate of *k*_− 1_. This 20-nt mode assumes constant *k*_1_. One scenario involves DBD-A, DBD-B, and DBD-C binding to ssDNA, with subsequent DBD-D binding leading to the 30-nt mode. In the 20-nt mode, DBD-D binds an extra 10-nt ssDNA with a rate of *k*_2_ and dissociates at *k*_− 2_, forming the 30-nt mode (FLBM). RPA always aligns in the same direction along DNA.

The multiple binding modes result in interesting kinetic features like facilitated exchange and desorption. In this model, the ssDNA fragment state is represented by a vector of length *L* with each component taking values in {0, 1}, where 0 indicates an unoccupied nucleotide and 1 indicates an occupied one. Each RPA initially occupies ℓ = 20 nts and can further occupy Δℓ = 10 nts if the local state allows, moving in the 3’ direction. Each DNA site can be occupied by only one RPA molecule, altering the RSA model’s available reactions. Each consecutive unoccupied segment of length ℓ recruits one RPA at the rate of *k*_1_, with the total binding rate given by:

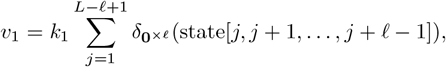

where *δ*_*a*_(*b*) is the Kronecker delta function, **0**^*×*ℓ^ is a zero vector of length ℓ, and state[*j*, …, *j* + ℓ 1] is the ssDNA state vector.

To occupy another 10 nts, we assigned a rate parameter *k*_2_ and calculate the overall rate by:

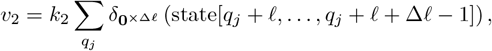

where *q*_*j*_ represents the leftmost position of each bound RPA in the 20-nt mode. For unbinding, the 30-nt mode reopens to the 20-nt mode at rate *k*_−2_, and the 20-nt mode desorbs at rate *k*_−1_. The overall rates are:

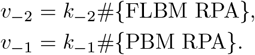

The total possible reaction rate is:

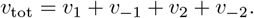

Reactions occur stochastically according to exponentially distributed waiting times with parameter *v*_tot_. The waiting time *δt* follows the exponential distribution with a probability density function:

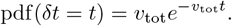

After each reaction, the DNA state updates, and the possible reactions are re-evaluated. The Gillespie algorithm was used to sample the trajectories of this stochastic model. We used Julia to perform exact stochastic simulations of all known RPA-ssDNA interactions, and codes are available at (https://github.com/hsianktin/RPA_model).

## Appendix C: Implementation of benchmarks

In this section, we introduce the benchmark methods; hyperparameters and settings for training of all methods used in this paper are shown in Table III.

**TABLE III.** Hyperparameters of benchmark methods.

| Model | Implementation |
| --- | --- |
| RNN | one RNN layer plus four MLP layers<br>width of RNN layer: 256<br>learning rate: $5 \times 10^{-4}$<br>learning epoch: 2000<br>Optimization method: Adam |
| LSTM | one LSTM layer plus four MLP layers<br>width of LSTM layer: 128<br>learning rate: $5 \times 10^{-4}$<br>learning epoch: 3000<br>Optimization method: Adam |

### 1. RNN networks

Recurrent neural networks (RNN) are often used for language processing, but they can also be used to analyze time series data with temporal correlations. For reconstructing the RPA-DNA binding dynamics, we used the RNN model in Ref. [66]. A neural network that contains two layers of RNN and two linear layers was used to model RPA’s dynamic binding with single-stranded DNA. All layers of RNN are built using `torch.nn.Module` package. Hyperparameters in the neural network is initialized by default. The parameters of this neural network are initialized by default. At each time step, ***X***(*t*; *ω*) ∈ ℝ^*d*^ representing the RPA dynamics at this time point is inputted in Example II B 2, and the dynamics at the next time point: ***X***(*t* + Δ*t*; *ω*) is outputted as the prediction. The RNN is trained by optimizing the loss function Eq. (8). The pipeline of the model could be seen in Fig. S1.

**Fig S1.**
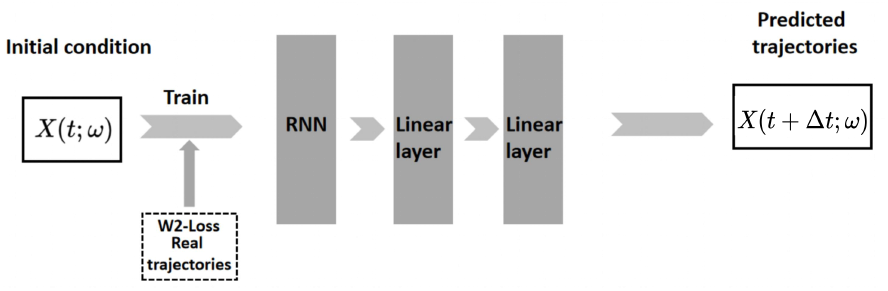
The structure of the RNN model used. The RNN layer in this figure can be built using `torch.nn.RNN`. The structure of RNN in the figure can be found in Ref. [67].

### 2. LSTM networks

Long Short-Term Memory (LSTM) networks [68], a class of variants of Recurrent Neural Networks (RNNs), have been widely used in modeling sequential data such as time series and natural language. We used the LSTM network model proposed in Ref. [69] implemented through the `torch.nn.Module` package for reconstructing the RPA-DNA dynamics. Hyperparameters in the neural network is initialized by default. At each time step, the ***X***(*t*; *ω*) ∈ ℝ^*d*^ discussed above is inputted in Example II B 2, and the state at the next time point: ***X***(*t* + Δ*t*; *ω*) is outputted as the prediction. Then, the gradients of the loss function Eq. (8) are calculated to update the parameters in the LSTM model. The input size of the two layers of LSTM is 2 and the hidden size is 4. The input size and output size of the two linear layers are 8,4 and 4,2 respectively. The LSTM is trained by optimizing the loss function Eq. (8). The pipeline of the model is shown in Fig. S2.

**Fig S2.**
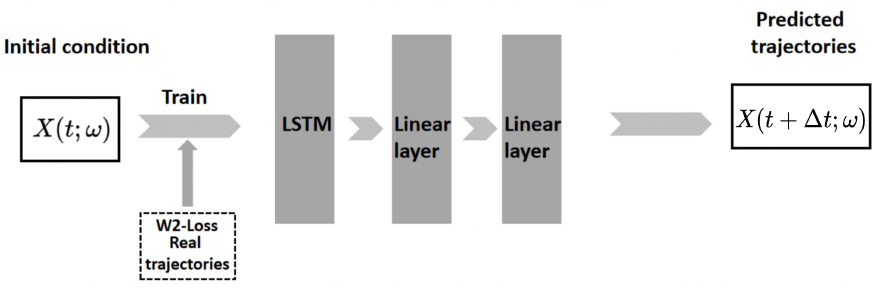
The structure of the LSTM model used. The LSTM layer in this figure can be built using `torch.nn.LSTM`. The structure of LSTM in figure can be found in [70].

### 3. Gaussian Process

The Gaussian Process (GP) [71] is also widely used in modeling time-series data; it is implemented using the `gaussianprocess.GaussianProcessRegressor` package in Python. We used radial basis functions as kernel functions (using sklearn.gaussianprocess.kernels package), and the hyperparameters of *α* and n-restarts-optimizer are 0.3 and 5, respectively. When training the GP model, we inputted the trajectories of all time points in the training set to the GP for fitting. Then, we inputted the trajectory of the current time point in the testing set, and the model predicts the trajectory of the next time point based on the kernel. The detailed structure of the GP model we used is the same as that in Ref. [72].

**Fig S3.**
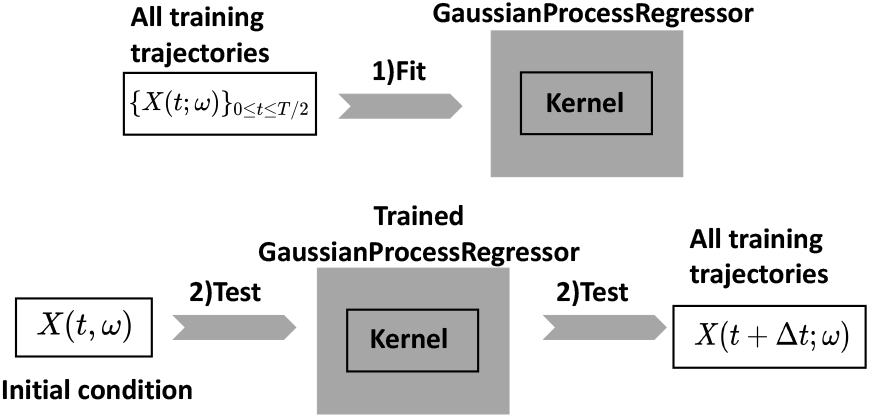
The structure of the GP model used in this work.

## Appendix D: Generating the simulated NF*κ*B dataset

To evaluate the performance of our proposed END-nSDE method in reconstructing dynamics of NF*κ*B signaling, we generated training trajectories by simulating a previously developed 52-dimensional model [46, 51] for the NF*κ*B signaling network under 100ng/mL TNF stimulation (see Eqs. (10) for the SDEs). The synthetic trajectories were generated using 100 sets of noise intensities. We set *σ*_1_ ∈ [10^−3.2^, 10^−2.2^], *σ*_2_ ∈ [10^−2.5^, 10^−1.5^] in Eqs. (10) and use 100 combinations 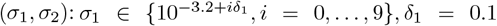 and *σ*_2_∈ 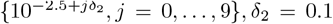 respectively. Other parameters were fixed constants. The parameter values in Eqs. (10) are listed in Table IV. The parameters for the remaining equations are the same as those in Refs. [46, 51].

**TABLE IV.** Parameter values for the NF*κ*B model.

| Parameter | Value | description |
| --- | --- | --- |
| $k_{\text{basal}}$ | $5 \times 10^{-7}$ | basal IκBα mRNA synthesis |
| $k_{\text{max}}$ | $6 \times 10^{-5}$ | maximal rate of IκBα mRNA synthesis induced by NFκB |
| $n_{\text{NF}\kappa\text{B}}$ | 2.938 | Hill coefficient for mRNA syn |
| $K_{\text{NF}\kappa\text{B}}$ | 0.1775 | EC50 for mRNA syn |
| $k_{\text{deg}}$ | 0.33 | degradation rate of IκBα mRNA |
| $k_{\text{imp}}$ | 0.6 | import rate of NFκB |
| $k_{\text{a-I}\kappa\text{B-NF}\kappa\text{B}}$ | 200 | association rate of NFκB and IκBα |
| $k_{\text{deg-NF}\kappa\text{B}}$ | 0 | degradation rate of NFκB |
| $k_{\text{exp}}$ | 0.042 | export rate of NFκB |
| $v$ | 3.5 | Volume ratio: cytoplasmic volume/nuclear volume |
| $k_{\text{d-I}\kappa\text{B-NF}\kappa\text{B}}$ | 0.008 | dissociation rate of NFκB-IκBα |
| $k_{\text{phos}}$ | 2 | Phosphorylation/degradation of complexed IκBα |

The corresponding SDEs were simulated using the ‘SDEProblem’ function from the ‘DifferentialEquations’ package in Julia. Simulations were conducted from 0 minutes (stimulus application time) to 150 minutes, and the state was recorded at every 5 minute intervals. Initial values were set to the steady-state solutions of the ordinary differential equations (ODEs), which were obtained using the ode15s function in MATLAB.

## Appendix E: Training a neural SDE using simulated NF*κ*B trajectories

We partitioned the simulated ground-truth trajectories into training and validation sets as follows: 50% of each of 96 sets of trajectories (out of a total of 121) associated with different noise intensities were used for training, while the remaining simulated ground-truth trajectories were used as the validation dataset. Since we could only observe NF*κ*B activity, we defined our loss function (see Eq. (8)) to focus solely on the nuclear NF*κ*B and the I*κ*B*α*-NF*κ*B nuclear complex. In the loss function Eq. (8), *ω*≡ (*σ*_1_, *σ*_2_) denoted two noise intensities (see Eqs. (10)). *µ*(*ω*) and 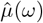 are the distributions of ***X***(*t*; *ω*) and 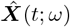,respectively. ***X***(*t*; *ω*) represented the values of (*u*_5_, *u*_10_) in Eqs. (10) at time *t*. The other 50 variables were not included in the calculation of the loss function.

## Appendix F: Reconstructing I*κ*B*α*NF*κ*Bn and NF*κ*Bn in NF*κ*B signaling as a 2D SDE model

To investigate the importance of latent variables that are not included in the loss function, instead of reconstructing the 52D surrogate SDE model for the NF*κ*B signaling dynamics, we attempted to directly reconstruct the dynamics of the nuclear complex I*κ*B*α* − NF*κ*B and nuclear NF*κ*B using the 2D SDE

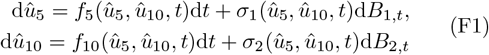

to approximate the two SDEs in NF*κ*B signaling model [46, 51] that describes nuclear I*κ*B*α*-NF*κ*B complex and NF*κ*B. Simulations and reconstruction of this model are shown in Fig. S4.

Using a 2D SDE model to reconstruct the NF*κ*B signaling dynamics could not accurately reconstruct the noisy dynamics of nuclear I*κ*B*α*-NF*κ*B and NF*κ*B. As shown in Figs. S4G-H, for certain noise strengths (*σ*_0_, *σ*_1_), both the training and testing losses (the temporally decoupled squared *W*_2_ distance Eq. (3)) are large compared to those obtained from the direct reconstruction of the full 52-dimensional model. Furthermore, when providing the inferred noise strengths from experimentally observed trajectories (the same as in Figs. 6A, B), the reconstructed 2D SDE fails to generate predicted trajectories (*û*_5_, *û*_10_) that align well with experimental data. Thus, it is necessary to retain the remaining 50 variables in the model, although they are not directly used in the calculation of the loss function.

## Appendix G: Dividing experimental NF*κ*B trajectories into subgroups

We divided the experimentally observed trajectories into 32 groups, each consisting of 32 trajectories. The experimental data were divided into different groups based on each trajectory’s correlation with a ODE-model-based deterministic trajectory [50, 51]

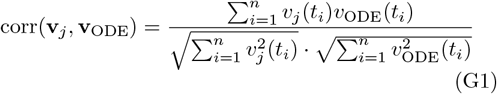

where **v**_*j*_(*t*_*i*_), **v**_ODE_(*t*_*i*_) denote the *j*^th^ observed trajectory in experimental data and the ODE trajectory. The closer a trajectory is to the first-principle-based ODE trajectories, the higher probability that the fluctuations result from intrinsic noise (*i*.*e*. the Brownian-type noise in Eq. (10)).

## Appendix H: Training the neural network to infer noise intensities in NF*κ*B dynamics

We trained a neural network that took a group trajectories as the input and then outputted inferred intrinsic noise intensity parameters (*σ*_1_ and *σ*_2_ in Eq. (10)) for the given group. Weights and biases in this neural network were optimized by minimizing the mean squared error (MSE) loss:

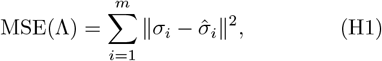

where *σi* are the ground truth noise intensity parameters underlying the *i*^th^ group of observed trajectories and 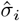 are the corresponding predicted parameters. Despite the assumption of all cells sharing the same drift function (the same underlying dynamics), different trajectories naturally arise from intrinsic noise.

**Fig S4.**
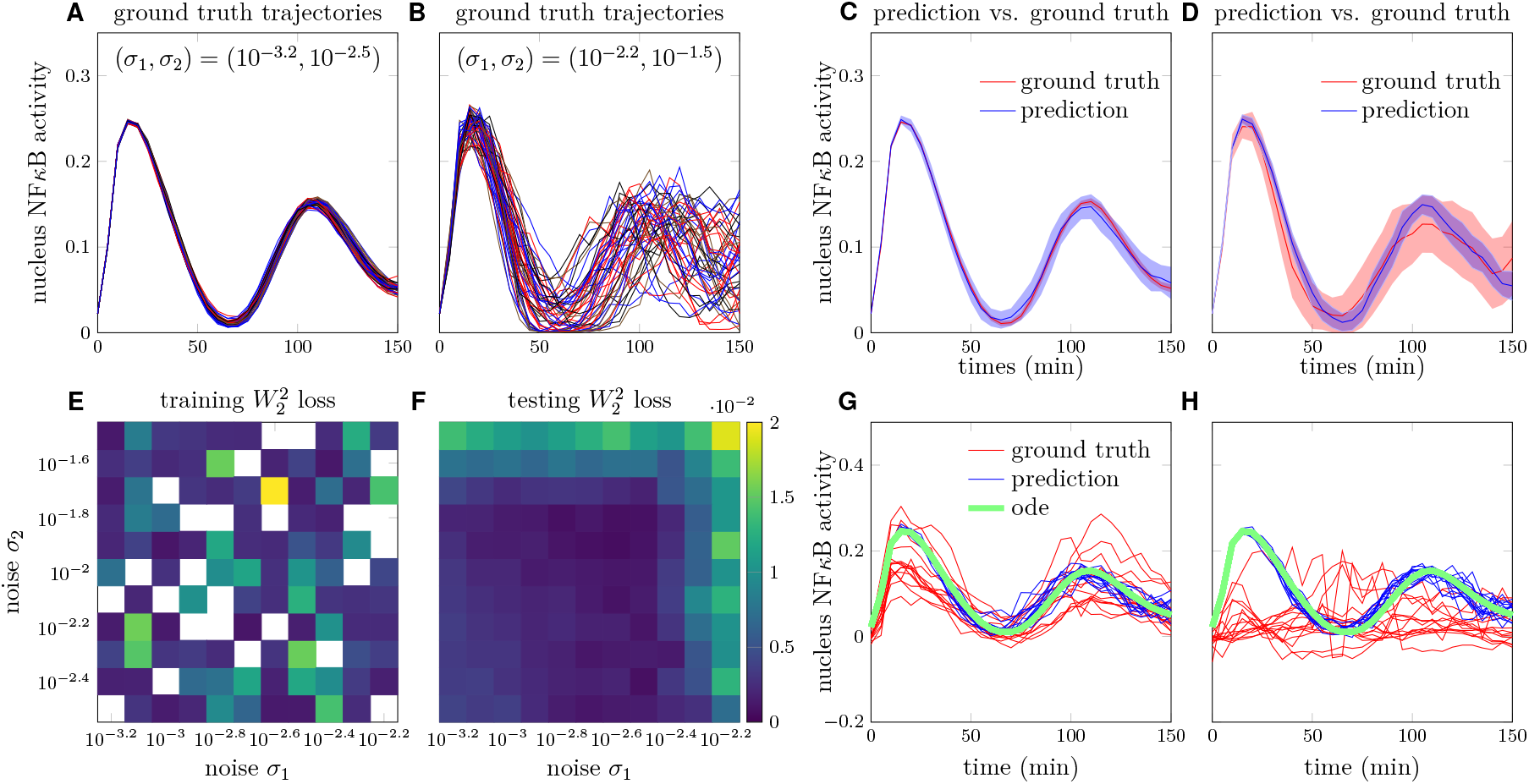
Reconstructed NF*κ*B dynamics using a two-dimensional nSDE. A. Trajectories of nuclear NF*κ*B concentration over time in the synthetic dataset with noise intensities *σ*_1_ = 10^−3.2^, *σ*_2_ = 10^−2.5^. B. trajectories of nuclear NF*κ*B concentration over time in the synthetic dataset with noise intensities *σ*_1_ = 10^−2.2^, *σ*_2_ = 10^−1.5^. C. Comparison of NF*κ*Bn trajectories predicted by the neural SDE with the ground truth trajectories under noise intensities *σ*_1_ = 10^−3.2^, *σ*_2_ = 10^−2.5^. D. Comparison of NF*κ*Bn predicted by the neural SDE with the ground truth trajectories under noise intensities *σ*_1_ = 10^−2.2^, *σ*_2_ = 10^−1.5^. E. The squared *W*_2_ distance between distributions of predicted trajectories and ground truth trajectories in the training set under different noise strengths *σ*_0_, *σ*_1_. Empty cells indicate that the corresponding parameter set is not included in the training set. F. The squared *W*_2_ distance between distributions of the predicted trajectories and the ground truth trajectories on the testing set. G.-H. After grouping the experimental trajectories by cosine similarity to the ODE reference trajectory (shown in green), the trained neural network estimated the noise (*σ*_1_, *σ*_2_) of each group of experimental trajectories (shown in red). Then, the estimated noise was fed into the nSDE to generate a group of (reconstructed) trajectories (shown in blue). The highest- and lowest-ranked similarity groups (#1 and #31, see Fig. 8D, E, I) are shown in G. and H., respectively.

We adopted a neural network that takes a group of trajectories (*u*_5_(*t*) + *u*_10_(*t*) in Eq. (10)) under the same noise intensity (*σ*_1_, *σ*_2_) as the input and outputs the inferred noise intensity 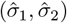.The training and testing sets are the same as those in Appendices D and E. The workflow for splitting NF*κ*B SDE-simulated trajectories into training and testing datasets is illustrated in Fig. S5A, B. Specifically, out of the 121 combinations of noise intensities (each containing 100 simulation trajectories), 20% were designated for the testing dataset pool. For the remaining 80% noise intensities, 50% of the trajectories under each parameter were randomly selected and added to the training dataset pool (indicated by the blue box in Fig. S5B). The remaining 50% of trajectories from these parameter sets were used for the testing dataset pool (red box in Fig. S5B). From both the training and testing dataset pools, for each combination of noise intensity, a group of trajectories were randomly sampled using a permutation sampling approach to construct the training and testing datasets (blue and red solid box in Fig. S5B).

The neural network used is equipped with an attention structure [73] followed by a feed-forward structure of 2 hidden layers with 64 and 128 neurons in each layer, respectively, where the attention mechanism is designed for assigning weights to different trajectories in a group (sequenced by their similarities to the deterministic ODE trajectory) (Fig.S5C). The hidden dimension of the attention structure in the query layer is 30 and the hidden dimensions of the attention structure in the key and value layers are both 32.

To assess the impact of group size on the performance of the trained neural network in predicting noise intensities, we tested group sizes corresponding to 2, 4, 8, 16, and 32 trajectories per group. The accuracy of the predictions was evaluated using the relative error metric:

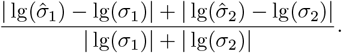

Detailed steps on inferring noise intensities from a group of trajectories are provided in Figs. S5A-C.

**Fig S5.**
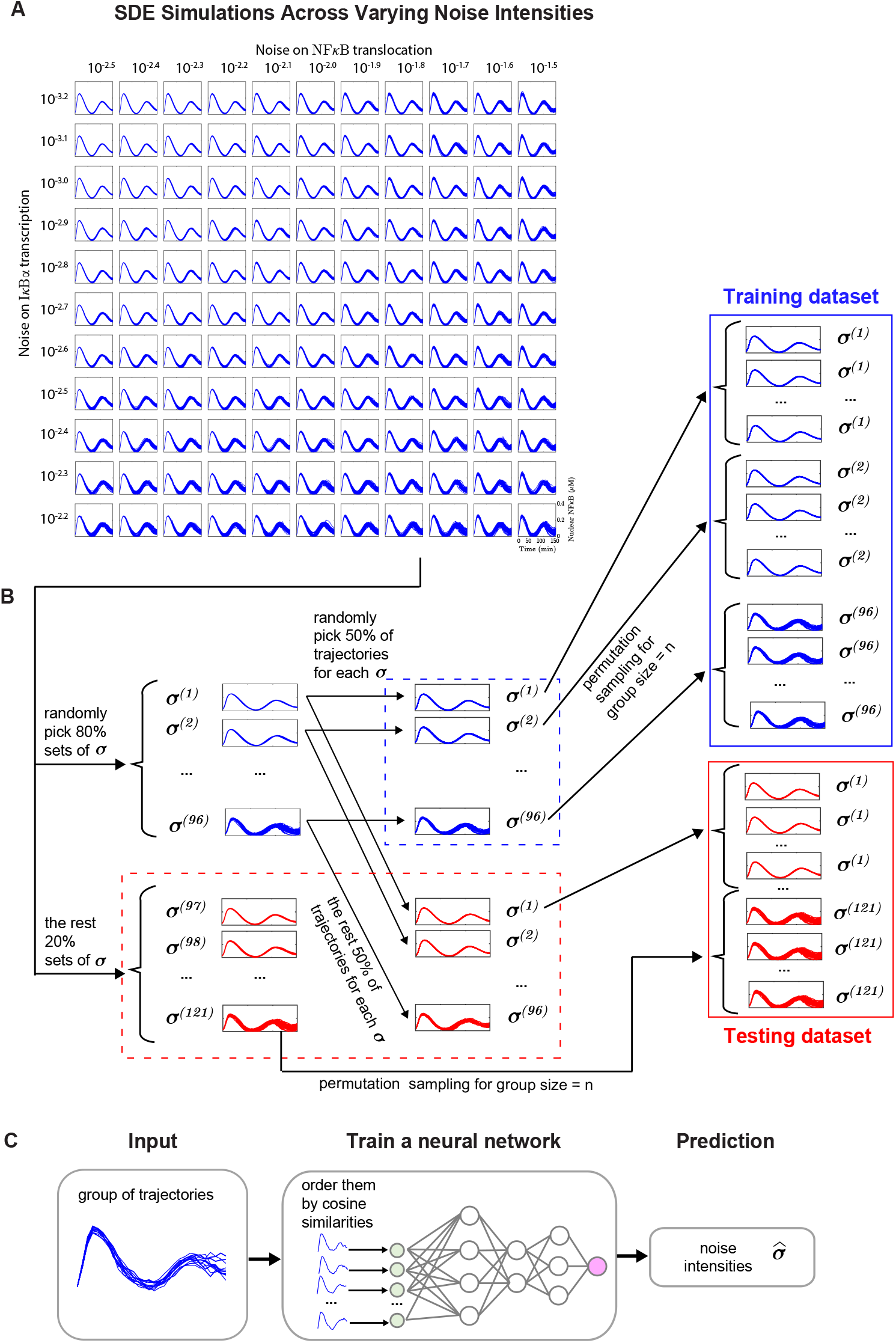
Workflow of using neural network to infer noise intensities from a group of trajectories. A. NF*κ*B SDE simulations under 121 different noise intensity settings. B. Schematic workflow for splitting the dataset into training and testing sets, where each group of trajectories is used to train and test the neural network for noise prediction. C. Schematic of the training of a neural network to predict noise intensities for each group of trajectories.

